# SALTClass: classifying clinical short notes using background knowledge from unlabeled data

**DOI:** 10.1101/801944

**Authors:** Ayoub Bagheri, Daniel Oberski, Arjan Sammani, Peter G.M. van der Heijden, Folkert W. Asselbergs

## Abstract

**Background:** With the increasing use of unstructured text in electronic health records, extracting useful related information has become a necessity. Text classification can be applied to extract patients’ medical history from clinical notes. However, the sparsity in clinical short notes, that is, excessively small word counts in the text, can lead to large classification errors. Previous studies demonstrated that natural language processing (NLP) can be useful in the text classification of clinical outcomes. We propose incorporating the knowledge from unlabeled data, as this may alleviate the problem of short noisy sparse text.

**Results:** The software package SALTClass (short and long text classifier) is a machine learning NLP toolkit. It uses seven clustering algorithms, namely, latent Dirichlet allocation, K-Means, MiniBatchK-Means, BIRCH, MeanShift, DBScan, and GMM. Smoothing methods are applied to the resulting cluster information to enrich the representation of sparse text. For the subsequent prediction step, SALTClass can be used on either the original document-term matrix or in an enrichment pipeline. To this end, ten different supervised classifiers have also been integrated into SALTClass. We demonstrate the effectiveness of the SALTClass NLP toolkit in the identification of patients’ family history in a Dutch clinical cardiovascular text corpus from University Medical Center Utrecht, the Netherlands.

**Conclusions:** The considerable amount of unstructured short text in healthcare applications, particularly in clinical cardiovascular notes, has created an urgent need for tools that can parse specific information from text reports. Using machine learning algorithms for enriching short text can improve the representation for further applications.

**Availability:** SALTClass can be downloaded as a Python package from Python Package Index (PyPI) website at https://pypi.org/project/saltclass and from GitHub at https://github.com/bagheria/saltclass.

## Background

A considerable amount of the data stored and documented in electronic health records (EHRs) are in the form of unstructured or semi-structured narrative text [1, 2, 3]. EHR text may contain short and noisy data. The telegraphic style of clinical narrative notes, including spelling errors or ambiguous abbreviations and measurements, renders information retrieval difficult [2, 3, 4]. In this context, traditional learning classifiers do not perform satisfactorily because clinical short notes do not provide sufficient word occurrences or shared context information, [5, 6]. By contrast, text mining and natural language processing (NLP) have become the most widely used big-data analytical techniques in healthcare applications [1].

Among the attempts to develop NLP software for healthcare [3, 7, 8, 9, 10, 11, 12, 13, 14, 15, 16], MedLEE (medical language extraction and encoding) [8], cTAKES (clinical text analysis and knowledge extraction system) [9], CogStack [10], and CLAMP [15] are prominent examples. MedLEE is an NLP system that can extract information from textual patient reports based on controlled vocabularies. It uses a lexicon to map terms into semantic classes, and a semantic grammar to generate formal representations of sentences. It then performs named entity recognition by dictionary look-up, handles abbreviations using a mapping table, and performs word sense disambiguation based on contextual rules. cTAKES is an NLP system for the extraction of information from electronic medical free text. It processes clinical text by identifying clinical entities such as drugs, diseases, and symptoms. CogStack implements data mining techniques that can search clinical data sources using automated information extraction of medical concepts. It uses NLP annotations to generate a timeline for patient interactions with services. CLAMP is abbreviated for clinical language annotation, modeling, and processing. It is a customizable NLP pipeline achieved good performance on named entity recognition and concept encoding for processing clinical text data. CLAMP’s components include sentence boundary detection, tokenizer, part-of-speech tagger, section header identification, abbreviation reorganization and disambiguation, named entity recognizer, and rule engine.

Sparsity in the feature space is a characteristic of clinical text that should be appropriately handled. To this end, studies on short-text data classification [5, 6, 17, 18, 19, 20, 21] have adopted two major approaches to enrich short text. One is to fetch contextual short-text information to add more text directly; this approach is called “dictionary-based method.” The other approach, which is called “topic-based method,” is to extract latent topics from an existing corpus. These topics are then used as features in further applications. The latter approach may lead to information loss, whereas the former cannot be applied in every domain owing to the lack of standard dictionaries, which is also a problem in EHR text classification.

Zelikovitz and Hirsh [22] presented a dictionary-based method to reduce error rates in short-text classification by using a large body of potentially unrelated background knowledge. They used an information-integration tool for text classification to query and integrate varied textual sources from the web. Dai et al. [18] proposed Crest, a topic-based method that generates topic clusters from training data. They used topic information to represent short texts by a new feature space. Crest represents short-text documents as augmented features using the cosine similarity between every document and the clusters. Cheng et al. [5] also proposed a topic-based method for short texts whereby topics are captured based on aggregated biterms in an entire corpus to tackle the sparsity problem. They define a biterm as an unordered word-pair co-occurring in a short text, considering the entire corpus a mixture of topics, where each biterm is drawn from a specific topic independently. Another study on short-text classification for medical records is the use of a bidirectional long short-term memory recurrent network by Cao et al. [23], where a knowledge-guided short-text classification system was proposed for healthcare applications using domain concept dictionaries. It was claimed that clinical notes contain domain-specific or infrequently appearing words. This may result in a poor embedding owing to the lack of training data.

Yu et al. [24] proposed LibShortText software, which is an open source tool for short text classification. LibShortText is a well-implemented Python package which provides a capability to change the parameters of the support vector machine algorithm. Yu et.al. have demonstrated that because short texts have more features than records the use of linear kernel for SVM algorithm is ideal.

In the present study, we develop a hybrid technique, which is called “intra-clustering approach,” that combines the advantages of both dictionary- and topic-based approaches. An important difference between the proposed technique and that in [23] is that the latter is a dictionary-based method in which if the domain knowledge dictionary is incomplete, a multi-task model is used to learn the domain knowledge dictionary jointly and perform the classification task, whereas the former completely relies on input data in an unsupervised manner. With the proposed intra-clustering approach, we use clustering algorithms to deploy new features from an internal knowledge acquisition scheme. A notable aspect of the proposed method is that it does not use categories of training examples to enrich the representation of short texts. However, a potential problem is that there are words in the test set that have not occurred in the training set. To overcome this, the proposed intra-clustering approach enhances the representation by incorporating background knowledge from unlabeled data. This allows new words in the test set to be incorporated in representation learning.

In what follows, we present SALTClass (short and long text classifier): a Python package that applies the proposed approach to clinical short text classification. We use SALTClass on a dataset collected by the Department of Cardiology of University Medical Center Utrecht (UMCU). In the experiments, the goal is to classify clinical sentences from patient notes and letters. The results from this classification can identify whether the clinical letters contain information about a patient’s medical history. This classification can be used in two ways: (i) to mine a patient’s history from clinical letters and present it to the EHR and (ii) as the first step for further standardization of clinical history (e.g., ICD10 and SNOMED).

## Implementation

Short-text classification can be defined as follows: given a set of documents with representation *D*, a label from a set of categories is assigned to each document. As short texts are also characterized by their representation sparsity, *D* should be optimized so that better performance may be achieved in the analysis of EHR text data.

Figure 1 shows the semantic flowchart of the SALTClass NLP toolkit. SALTClass optimizes the representation *D* for short-text classification. This toolkit classifies sentences by cleaning them and using a combination of clustering algorithms and supervised classifiers. Let *D* and *D*^∗^ be the original clinical free-text dataset and the enriched dataset, respectively. In the framework of SALTClass, *D*^∗^ is enriched based on related knowledge from word clusters extracted by unsupervised clustering algorithms.

**Figure 1.**
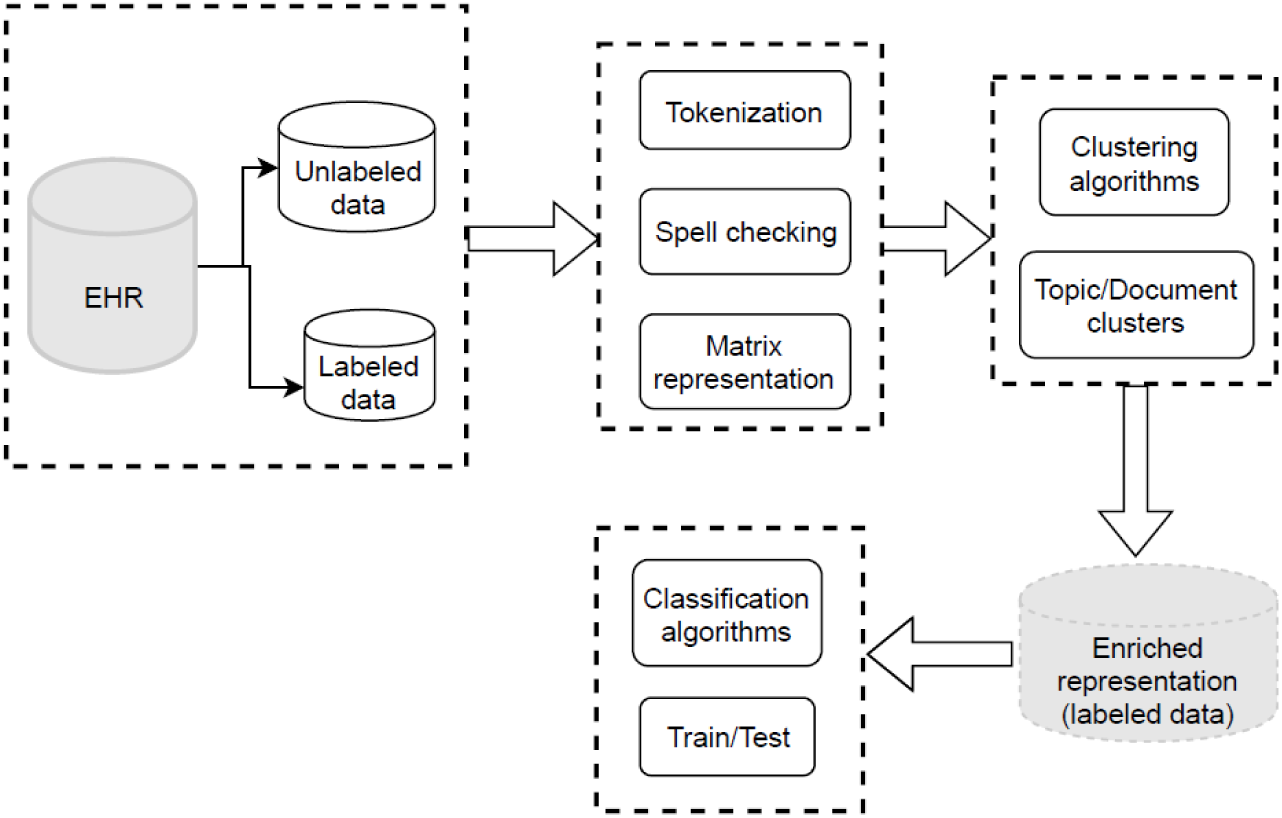
Architecture of SALTClass NLP software.

The first step of the SALTClass architecture is preprocessing, which includes detecting and extracting all sentences from *D*. Sentences will then be split using a tokenization module and will be represented by a vector-space bag-of-words model. The preprocessing step also includes removing spelling errors and stop words from *D*. The final two steps in the SALTClass architecture are the use of an unsupervised clustering procedure and a supervised classification algorithm.

In SALTClass, the unsupervised intra-clustering procedure is the heart of the architecture. This procedure uses the background knowledge in the data to optimize the vector representation. It pumps cluster information throughout the text body using a smoothing technique. This procedure provides text fragments with additional length. Intra-clustering is a hybrid technique, using the advantages of different modules, including dictionary- and topic-based approaches, smoothing methods, and cluster information. Dictionary-based techniques integrate text data with meta-information from other information sources, such as Wikipedia and WordNet, and topic-based methods represent short text with latent topics from the dataset. The intra-clustering algorithm in the proposed NLP toolkit is presented in Figure 2.

**Figure 2.**
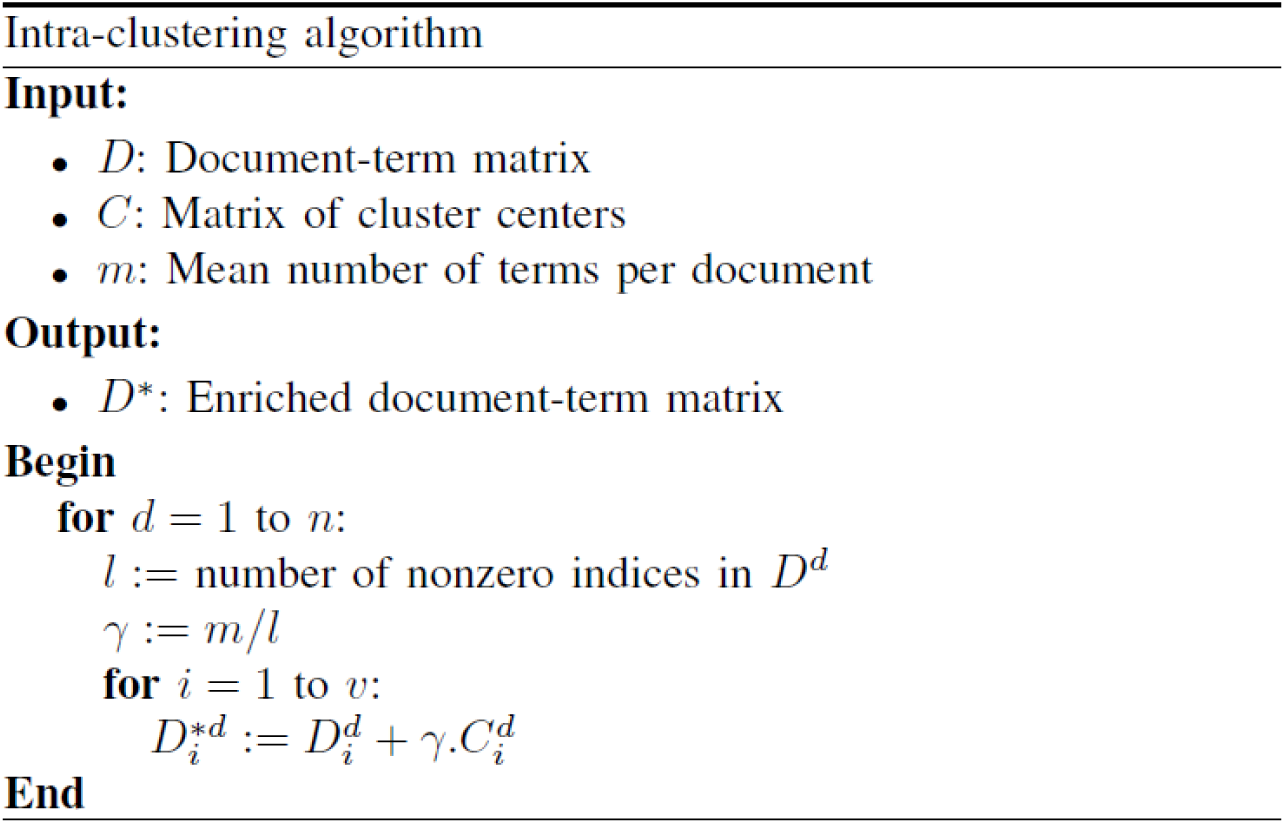
Proposed intra-clustering algorithm.

The intra-clustering algorithm expects three objects as input: *D*, *C*, and *m*. *D* is the document-term matrix for the text dataset. In this matrix, documents are represented by rows and terms (n-gram words) by columns, and the elements are the counts or the weights. It is not necessary to provide labeled categories with the matrix *D*; hence, the intra-clustering algorithm uses topic distributions as the background knowledge of the entire dataset.

*C* is the matrix of cluster centers, which is the cluster-term matrix. *m* is the average number of terms per document in the dataset. *n* is the total number of documents. 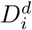 denotes term *i* in document *d* of dataset *D*. *v* is the vocabulary size, that is, the number of unique terms in the dataset. *l* counts the number of individual terms in document *d* for calculating its enrichment weight *γ*. The intra-clustering algorithm outputs the enriched representation in the document-term matrix *D*^∗^.

Figure 3 shows an illustrative example of the intra-clustering algorithm. As can be seen from this example, the proposed intra-clustering method attempts to move documents toward the center of assigned clusters.

**Figure 3.**
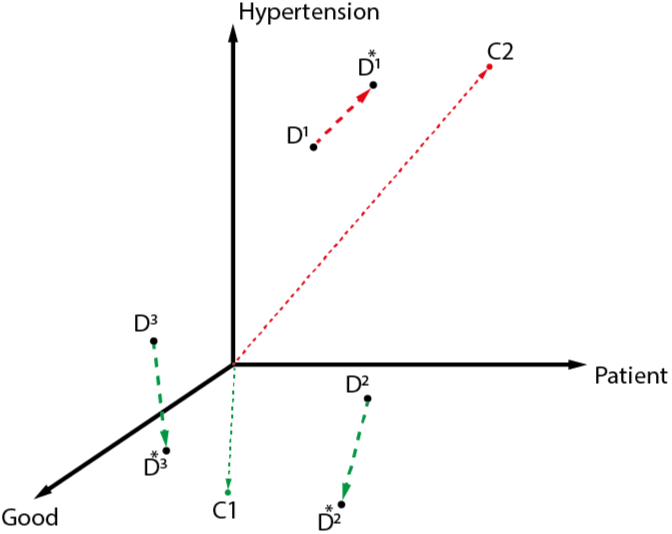
Illustrative example of the proposed intra-clustering algorithm. The feature space in this example has three words: Patient, Hypertension, Good. *D*, *C*, and *D\** denote the original document, the cluster center, and the enriched document, respectively. The documents are enriched toward the center of the cluster to which they belong.

This procedure may add words from the center vector to the document vector and change the values of the document vector representation. That is, the proposed algorithm represents short texts by adding information from latent topics to the original annotated dataset. This algorithm is an unsupervised learner, and instead of using training data, it uses both training and test examples in choosing the hypothesis of the learner. For classifying clinical short texts, the proposed toolkit allows the unseen words from the test set to be incorporated in the process of representation learning.

With SALTClass, different methods are implemented to enable users to choose a configuration. A summary of the main methods and parameters in SALTClass is shown in Tables 1 and 2, respectively.

**Table 1.** Methods overview

| Method | Description | Parameters |
| --- | --- | --- |
| SALT | To initialize an object with a training matrix and define a setting | x, y, vocabulary, kwargs |
| SALT.data_from_dir | To initialize an object from a directory of text files | train_dir, kwargs |
| initialize_dataset | To read documents from a training path and vectorize them | train_dir, word_vectorizer, language |
| enrich | To enrich a training set with a method | method, num_clusters, include_unlabeled, unlabeled_dir, unlabeled_matrix |
| train | To train a supervised classifier | kwargs |
| predict | To predict a category | data_file |

**Table 2.** Parameters definition

| Parameters | definition |
| --- | --- |
| x | Training data: a numerical document-term matrix |
| y | Target values: categories |
| vocabulary | Variables (word features) in x |
| kwargs (SALT method) | Keyword arguments:<br>language='nl',<br>vectorizer='count' |
| train_dir | Training data: directory folders of text files |
| kwargs (SALT.data_from_dir method) | Keyword arguments:<br>language='nl',<br>vectorizer='count' |
| word_vectorizer | Word feature vectorizer: 'count', 'tfidf' |
| language | Language of text: 'nl', 'en' |
| method | Clustering method: lda, kmeans, mbk, birch, gmm, ms, dbscan |
| num_clusters | Number of clusters |
| include_unlabeled | Flag: True, False |
| unlabeled_dir | Directory folder of unlabeled text files |
| unlabeled_matrix | Unlabeled document-term matrix |
| kwargs (training method) | Keyword arguments:<br>classifier='SVM', kernel='poly', degree=2<br>classifier='SVM', kernel='sigmoid'<br>classifier='SVM', kernel='lin', gamma=2<br>classifier='KNN', k=3<br>classifier='DT', depth=5<br>classifier='RF', depth=5, n_estimators=10, max_features=1<br>classifier='NN', alpha=1, hidden_layer_sizes=(50,), max_iter=10,<br>solver='sgd', 'adam', activation='relu'<br>classifier='AdaB'<br>classifier='GaussianNB'<br>classifier='MultinomialNB'<br>classifier='GP'<br>classifier='LR' |
| data_file | A text file for prediction |

The list of clustering and classification methods implemented in SALTClass are:

- K-Means
- MiniBatch K-Means
- Latent Dirichlet allocation (LDA)
- Balanced iterative reducing and clustering using hierarchies (BIRCH)
- Density-based spatial clustering of applications with noise (DBSCAN)
- Gaussian-mixture modeling (GMM)
- Meanshift
- Logistic regression (LR)
- Neural networks (NNs)
- K-Nearest neighbors (KNNs)
- Support vector machines (SVMs)
- Decision trees (DTs)
- Random forest (RF)
- AdaBoost (AdaB)
- Gaussian naive Bayes (GaussianNB)
- Multinomial naive Bayes (MultinomialNB)
- Gaussian processes (GPs)

## Experiment study

### Example

We present an example to clarify the principle of SALTCLass. In this example, we employ the K-Means and LDA clustering methods from the SALTClass package to illustrate the concept of enrichment. The example has a dataset of five sentences (short documents), which are shown in Figure 4. This figure also shows the count-vector representations for the short documents. As the vocabulary is small, we do not remove stop words. The average number of terms in the documents is *m* = 4.4. The enrichment weight *γ* is also shown in Figure 4 and is calculated for each document using *m* and the inverse document length.

**Figure 4.**
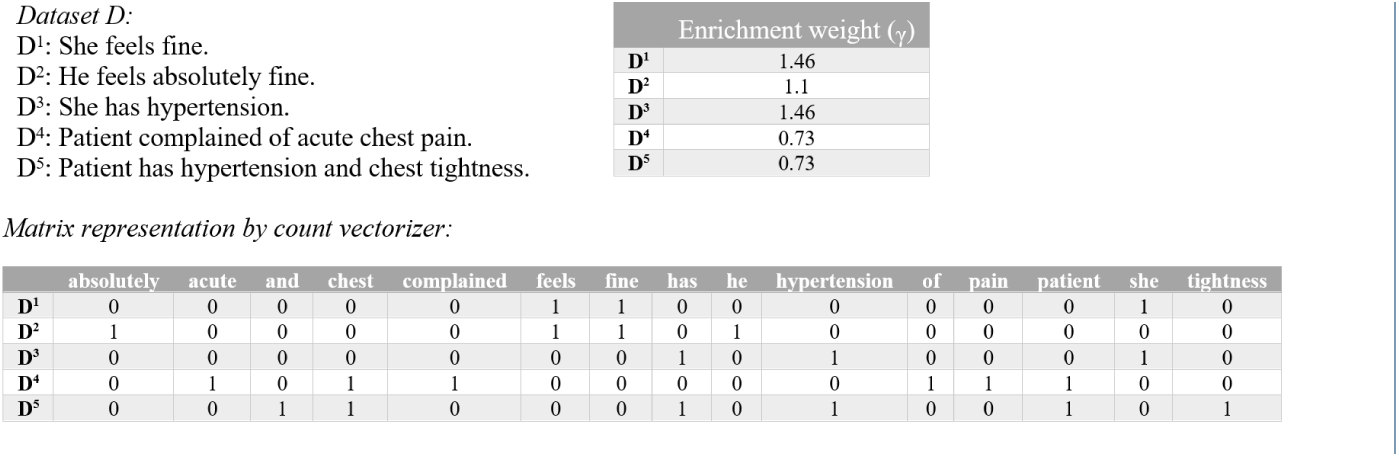
Example of enrichment representation. The data are represented with CountVectorizer from the Scikit-learn package in Python.

Figures 5 and 6 show the enrichment phase using the K-Means and LDA algorithms, respectively. For intra-clustering, we use the algorithms to generate two document clusters. This can be accomplished in the SALTClass package by using the command *object.enrich*(*method* =′ *kmeans*′, *num clusters* = 2) for the K-Means algorithm. For LDA, we use *object.enrich*(*method* =′ *lda*′, *num clusters* = 2). The intra-clustering algorithm first calculates the enrichment weight *γ* for each document. Subsequently, the count vectors are enriched using the clustering outputs. In the K-Means algorithm, the count vectors use the corresponding center vector from the clusters to update their values. In the LDA algorithm, the count vectors use both the document topic and the topic word distributions to modify the representation.

**Figure 5.**
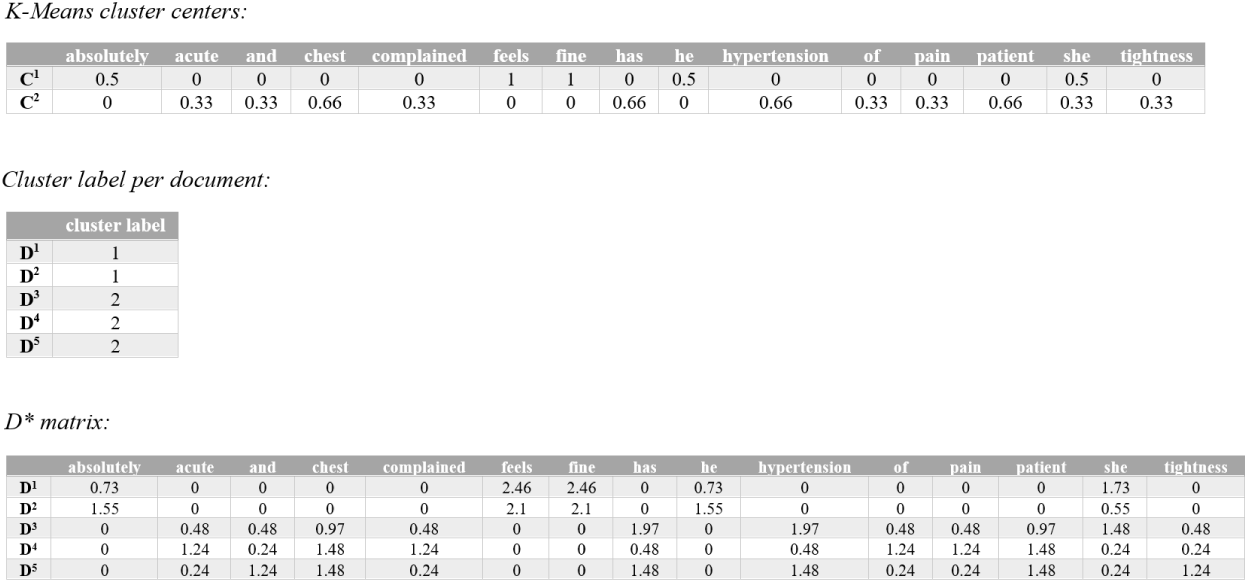
Intra-clustering with K-Means. The K-Means clustering method is applied to the example to generate two document clusters. The matrix *D\** contains the enriched representation for the documents.

**Figure 6.**
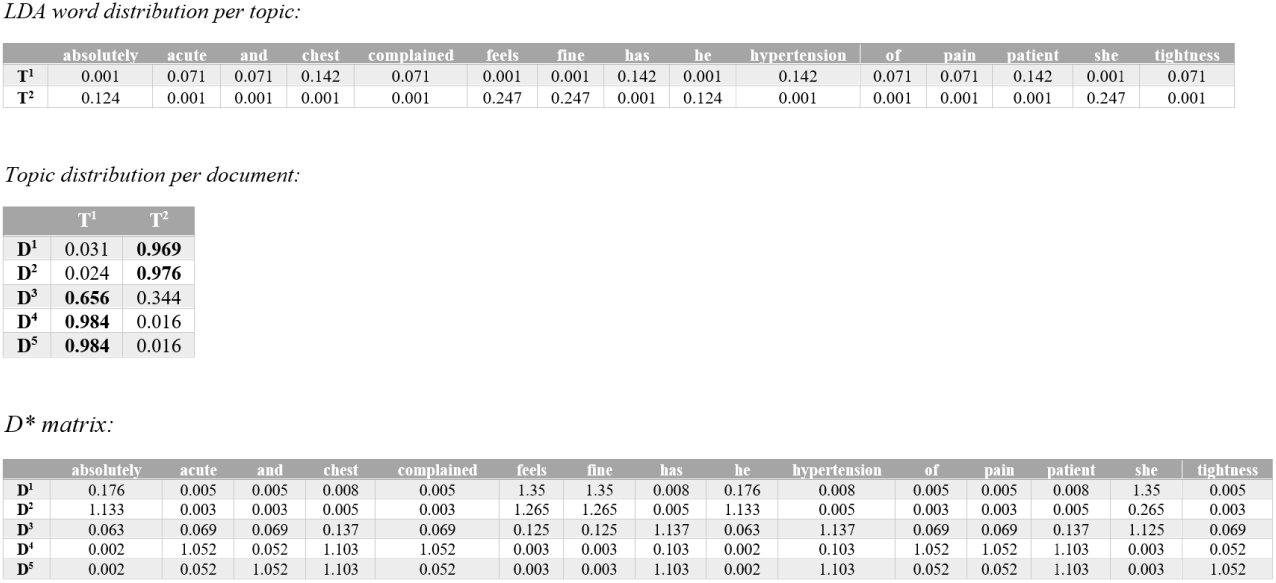
Intra-clustering with LDA topic modeling. The LDA algorithm is applied to the example to generate two document clusters. The matrix *D\** contains the enriched representation for the documents.

Figure 5 also shows the K-Means clustering labels for each document. Figure 6 shows the word distributions per topic and the topic distributions per document. It can be seen that the algorithm outputs the matrix *D*^∗^. K-Means and LDA clustering generate two different *D*^∗^ matrices. One noticeable difference between the outputs is the zero and non-zero values for the same cell. This is due to the LDA assumption that documents have multiple topics [25].

### Data

UMCU is one of the largest university hospitals in the Netherlands that provide specialized cardiac care. Given the structure of their EHRs, data are available in a research data platform and can be extracted accordingly. The textual dataset used in this study comprises all clinical cardiovascular notes from medical doctors or physician assistants between 2014 and 2018. 1002 Dutch clinical notes have been manually annotated for medical history, based on the international classification of disease (ICD10) criteria and were sample-wise checked by medical doctors. Conflicts in annotation were resolved through discussion. The words in these clinical notes on which the annotation was based were also marked for text mining purposes. These words delineate sentences that contain medical history. The training data set was generated based on this delineation. After annotation, the training data for short text classification contained 11,053 sentences, where 3,560 of them were related to medical history. Along with the labeled clinical notes, 20,200 unlabeled clinical cardiovascular sentences were used for the experimental study of the proposed NLP software.

#### Dutch text preprocessing

In NLP, a major type of preprocessing is to filter out stop words and spelling errors. Clinical notes may have misspellings and useless words, which are referred to as stop-words. These can be removed without any negative consequences to the training model. There is no universal list of stop words because a word can be meaningful or meaningless depending on the context. Making a list of stop words for an EHR is beyond the scope of this study. However, the NLTK ^[1]^ Python package provides lists of stop words in 21 different languages, including a list of Dutch stop words. To handle spelling errors in texts, we used the Python package language-check ^[2]^, which is a wrapper for the LanguageTool ^[3]^ package. LanguageTool is an open source proofreading software that can detect and correct spelling errors in more than 20 different languages. This package is freely available under the LGPL 2.1 or later. LanguageTool functions properly with Python 3.6 and JDK 8.

## Results

We use precision, recall and F1 score results by 5-fold cross validation in the experiments to evaluate the performance of the SALTClass toolkit. Precision is defined as the fraction of relevant documents among the retrieved documents, whereas recall is the fraction of relevant documents that have been retrieved over the total amount of relevant documents. The F1 score is defined as the harmonic average of the precision and recall of the test [26].

Table 3 shows the F1 score results of the first experiment, where we checked the effectiveness of SALTClass using *count* (*term frequency*) and *tfidf* vector representations. Setting1 is the count-vector representation and Setting2 is the *tfidf* representation. *tfidf* is short for “term frequency inverse document frequency” and is a numerical statistic that is intended to reflect the importance of a word to a document in a collection of documents. Neither Setting1 nor Setting2 calls the *enrich* function. These two settings are coupled in four different experiments with the following supervised classifiers:

**Table 3.**
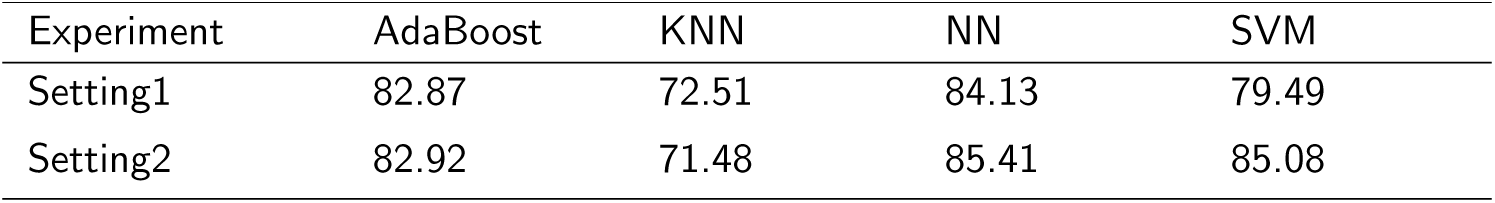
F1 score results of two representations with the learning classifiers

| Experiment | AdaBoost | KNN | NN | SVM |
| --- | --- | --- | --- | --- |
| Setting1 | 82.87 | 72.51 | 84.13 | 79.49 |
| Setting2 | 82.92 | 71.48 | 85.41 | 85.08 |

- **AdaBoost** is a type of ensemble learning for classification.
- **KNN** is a non-parametric instance-based learning algorithm.
- **NN** learns to map the input data to the output labels through a series of nonlinear compositions.
- **SVM** learns an objective function by employing internally a kernel trick.

As shown in Table 3, the results for the classifiers are highly similar in both representations. SVM and NN obtained much improved results with *tfidf*. KNN performed better with count representation, and AdaBoost gaind similar results with two representations. Nevertheless, the results of the two bag-of-words models demonstrate the average superiority of *tfidf* over count-vector representation. This is because *tfidf* extracts more descriptive features from a document.

Tables 4 and 5 compare the precision and recall results for three settings, respectively. Setting2 is the *tfidf* representation denoted by *Raw*. Setting3 is the *tfidf* representation calling the *enrich(num_clusters=10)* function to apply the intra-clustering algorithm. This experiment shows the results of the K-Means and LDA algorithms, as the unsupervised method used in the framework of the intra-clustering algorithm. We chose 10 as the number of clusters based on the experiments on different values. Setting4 is the same as Setting3 assigning *True* to the parameter *include_unlabeled* and a directory path to the folder of unlabeled data in the parameter *unlabeled_dir*.

**Table 4.** Precision results of the learning classifiers

| Method | Experiment | AdaBoost | KNN | NN | SVM |
| --- | --- | --- | --- | --- | --- |
| Raw | Setting2 | 79.64 | 70.88 | 82.56 | 82.21 |
| K-Means | Setting3 | 80.46 | 71.17 | 82.67 | 81.99 |
|  | Setting4 | 80.52 | 71.22 | 82.26 | 82.11 |
| LDA | Setting3 | 82.97 | 71.32 | 85.14 | 83.65 |
|  | Setting4 | 82.41 | 71.06 | 85.91 | 82.78 |

**Table 5.** Recall results of the learning classifiers

| Method | Experiment | AdaBoost | KNN | NN | SVM |
| --- | --- | --- | --- | --- | --- |
| Raw | Setting2 | 86.48 | 72.10 | 88.47 | 88.16 |
| K-Means | Setting3 | 86.56 | 72.11 | 88.13 | 88.19 |
|  | Setting4 | 86.73 | 72.47 | 88.74 | 88.42 |
| LDA | Setting3 | 88.49 | 73.16 | 89.72 | 89.82 |
|  | Setting4 | 88.69 | 72.84 | 92.71 | 91.49 |

The precision results in Table 4 demonstrate the effectiveness of Setting3 (using intra-clustering) and Setting4 compared with Setting2 with respect to medical history classification.

The highest precision of the intra-clustering algorithm was 85.91%, occurred when the NN classifier was used with unlabeled data. Comparing classifiers, the intra-clustering algorithm using NN has the highest improvement over the Raw representation. In this experiment, enriching the dataset improved the results for the four classifiers, when the LDA method was used as the clustering algorithm. However, the intra-clustering using the K-Means method improves the results in AdaBoost, KNN, and the Setting3 of NN. The intra-clustering using K-Means with the SVM classifier with the *tfidf* representation in Setting2 had slightly better performance than SVM in Setting3 and Setting4.

Table 5 shows the recall results of the proposed toolkit in different settings. These results show the improvement in the performance of the proposed approach. It is remarkable that the results for recall were higher than those for precision in all classifiers.

The results in Tables 4 and 5 show the superiority of using unlabeled data in the framework of the intra-clustering algorithm. The highest improvements were with the NN classifier in precision (0.77%) and recall (2.99%). Thus, when more data for clinical text classification are available, the proposed method has a higher chance of improving performance.

### Comparison with LibShortText software

LibShortText is a modification of the widely used LibSVM library, but for short text classification. The package includes the source code in Python 2.6 and C/C++. The LibShortText software implements document search using the vector space model, then uses a bag-of-word model to generate future vectors. The LibShortText software uses the LibSVM with the advantage of not being required to tune the algorithm for the optimum kernel function and penalty factor.

Table 6 shows the comparison of classification performance using LibShortText and SALTClass for the UMCU text data in terms of macro-precision, recall and F1 score.

**Table 6.** Results for the SALTClass and the LibShortText software

| Software | Precision | Recall | F1 |
| --- | --- | --- | --- |
| LibShortText | 81.95 | 87.66 | 84.61 |
| SALTClass | 85.91 | 92.71 | 89.11 |

For given short texts, LibShortText follows the bag-of-word model to generate features, and preprocess short texts by tokenization, stemming, and stop-word removal. The library also allows users to choose between unigram and bigram features. However, LibShortText follows the routine pipeline of text classification using the LibSVM library. On the contrary, SALTClass enriches the representation of short texts to improve the classification performance. As shown in Table 6, the SALTClass package gained better results compared with those on UMCU data from the LibShortText software.

## Conclusion

EHRs contain a wealth of information in clinical text form. Thus, text-mining techniques may be advantageously used to extract structured information. Medical data suffer from insufficient context, as they are represented in short texts. The SALTClass software was proposed to mitigate the classification error inherent in short texts by interpolating between observed and fitted counts obtained by clustering algorithm. SALTClass is a Python module for short-text classification built under the MIT license. This module contains several functions for text preprocessing, cleaning, clustering, and classification. With the proposed intra-clustering algorithm, SALTClass can be fed with unlabeled data to incorporate the background knowledge for short texts. The SALTClass NLP toolkit enables users to apply various configuration combinations to their case study. To evaluate the effectiveness of SALTClass, we analyzed the classification of short cardiovascular notes collected in the UMCU hospital. It was demonstrated that using SALTClass can improve classification performance in terms of the precision, recall and F1 score.

## Competing interests

The authors declare that the research was conducted in the absence of any commercial or financial relationships that could be construed as a potential conflict of interest.

## Author’s contributions

AB and DO designed the framework. AB wrote the code. All authors analyzed the data and experiments. All authors wrote the paper. All authors read and approved the final manuscript.

## Acknowledgements

F.W. Asselbergs is supported by UCL Hospitals NIHR Biomedical Research Centre.

## Ethics approval and consent to participate

Not applicable

## Consent for publication

Not applicable

## Footnotes

[1] https://www.nltk.org/

[2] https://github.com/myint/language-check

[3] https://languagetool.org

